# Embedding the de Bruijn graph, and applications to metagenomics

**DOI:** 10.1101/2020.03.06.980979

**Authors:** Romain Menegaux, Jean-Philippe Vert

## Abstract

Fast mapping of sequencing reads to taxonomic clades is a crucial step in metagenomics, which however raises computational challenges as the numbers of reads and of taxonomic clades increases. Besides alignment-based methods, which are accurate but computational costly, faster compositional approaches have recently been proposed to predict the taxonomic clade of a read based on the set of *k*-mers it contains. Machine learning-based compositional approaches, in particular, have recently reached accuracies similar to alignment-based models, while being considerably faster. It has been observed that the accuracy of these models increases with the length *k* of the *k*-mers they use, however existing methods are limited to handle *k*-mers of lengths up to *k* = 12 or 13 because of their large memory footprint needed to store the model coefficients for each possible *k*-mer. In order to explore the performance of machine learning-based compositional approaches for longer *k*-mers than currently possible, we propose to reduce the memory footprint of these methods by binning together *k*-mers that appear together in the sequencing reads used to train the models. We achieve this binning by learning a vector embedding for the vertices of a compacted de Bruijn graph, allowing us to embed any DNA sequence in a low-dimensional vector space where a machine learning system can be trained. The resulting method, which we call `Brume`, allows us to train compositional machine learning-based models with *k*-mers of length up to *k* = 31. We show on two metagenomics benchmark that `Brume` reaches better performance than previously achieved, thanks to the use of longer *k*-mers.

## 1 Introduction

The cost of DNA sequencing has reduced dramatically over the past years. Among its many applications, it has become the method of choice for metagenomics, a field that aims at characterizing an environment directly from the DNA it contains by sequencing DNA samples randomly collected from the environment. A bottleneck in modern metagenomics pipelines is the taxonomic binning step, i.e., the mapping of the output of second generation shotgun sequencing machines – millions to billions of short DNA reads – to known taxonomic clades. Of the several methods developed to tackle this problem, a natural one is to use an all-purpose DNA aligner, such as Li (2013), Langmead *et al.* (2009) or Li (2018), to align the reads to reference genomes. These aligners, which are usually based on string-matching algorithms, are very accurate but typically computationally too slow for taxonomic binning. More recently, competitive performances have been achieved by so-called compositional approaches, in terms of both speed and precision. Compositional approaches do not try to align a candidate read to a reference genome, but instead directly predict the genome the read is likely to come from based on the composition of the read in shorter strings, called *k*-mers. These methods can be roughly separated in two separate classes. Methods of the first group, so-called pseudo-alignment methods (Wood and Salzberg, 2014; Wood *et al.*, 2019; Ounit *et al.*, 2015), break a read into long and hopefully characteristic subsequences (*k*-mers, *k* ∼ 30) and match the read to the genomes/species that has the most *k*-mer matches. Usually a single match on those long *k*-mers is enough to ensure classification to a taxonomic clade. The second class of compositional methods treat the problem as a machine learning classification problem. Most of these approaches represent the DNA read as a set of short *k*-mers (*k* from 4 to 14) and then use a discriminative machine learning model to predict the clade of a read from the vector of frequencies of all *k*-mers it contains. In Vervier *et al.* (2016), this model is simply linear, akin to the bag-of-words model in natural language processing, and *k*-mers of lengths up to *k* = 12 are considered. Menegaux and Vert (2019) use a two-layer model called `fastDNA`, which first embeds the *k*-mers in a continuous low-dimensional vector space and then classifies reads in the vector space with multinomial logistic regression. The low-dimensional embedding in Menegaux and Vert (2019) allows them to reduce the memory footprint of the model when many possible taxonomic clades exist, and to consider *k*-mers of length up to *k* = 13. More recently deep learning has also been applied; GeNet encodes individual nucleotides (i.e., *k* = 1) followed by a convolutional neural network (Rojas-Carulla *et al.*, 2019), while DeepMicrobes shows the benefits of first encoding longer *k*-mers as input to a convolutional or recurrent deep learning architecture to boost the accuracy of the model (Liang *et al.*, 2019); they show that performance increases with *k* up to *k* = 12, the largest possible that allows to fit the model in the memory of the hardware needed to train the model. Georgiou *et al.* (2019) manage to reduce the memory footprint of the model by using a locality-sensitive hashing (LSH) of *k*-mers, allowing them to test models up to *k* = 13.

Results from Vervier *et al.* (2016); Menegaux and Vert (2019); Liang *et al.* (2019) all suggest that *k*-mers with larger *k* lead to better performance. Storing the parameters of the models in memory is however often the limiting factor to increase *k*, since the memory footprint of the model is typically *O*(*c*4^*k*^) when a model is stored for each of *c* clades (Vervier *et al.*, 2016), or *O*(*d*4^*k*^) when the *k*-mers are first represented in a *d*-dimensional vector space (Menegaux and Vert, 2019; Liang *et al.*, 2019). The exponential growth of the memory footprint with *k* has thus far limited the use of machine learning-based methods to *k* = 12 or 13. Hashing-based indexing of *k*-mers (Vervier *et al.*, 2016; Georgiou *et al.*, 2019) allow technically to increase *k* while keeping the memory footprint constrained, but the collisions created by hashing are detrimental to the performance of the model, resulting in practice to performance loss when *k* is larger than 13. Pseudo-alignment methods, on the other hand, work with longer discriminative *k*-mers (*k* between 20 and 31), but then use a very simple presence/absence model to classify the reads, and do not need to store any parameter for all possible *k*-mers.

In this work, we propose a new approach to reduce of memory footprint of machine learning-based models for large *k*, which allows us to explore the performance of these models for *k* up to 31. Our approach is based on binning together *k*-mers which occur consecutively in the same reads in the training set, and learning a single vector representation for each such bin. These bins, also called contigs or unitigs, correspond to vertices in a compact de Bruijn graph of the training sequences. Instead of storing vector representations for 4^*k*^ *k*-mers, our approach thus reduces the need to store representations for only *n*_*c*_ contigs, which can be much smaller than 4^*k*^.

We implement this idea in a software, `Brume`, which extends the two-layer model of Menegaux and Vert (2019) by embedding contigs instead of *k*-mers in the first layer. We report promising results of `Brume` on two metagenomics benchmarks, where increasing *k* beyond 12 or 13 leads not only to better performance compared to smaller *k*, but also compared to alignment-based approaches.

## 2 Approach

### 2.1 Two-layer classifier

Here we describe the two-layer machine learning model for read classification deployed in fastDNA (Menegaux and Vert, 2019), which we generalize below. We consider a finite alphabet 𝒜, which for our purpose is 𝒜 = {`A, C, G, T`}, the set of four nucleotides found in DNA. A *k*-mer *x* ∈ 𝒜^*k*^ is a string of fixed length *k*. In `fastDNA`, a *d*-dimensional embedding is first learned for each *k*-mer. If *N* = 4^*k*^ is the total number of possible *k*-mers, the embeddings matrix *M* is a *N* × *d* matrix with each row *M*_*s*_ coding the vector representation of a *k*-mer *s*. From the embedding matrix *M* of all *k*-mers, we then define the embedding Φ^*M*^ (**x**) ∈ ℝ^*d*^ of a read **x** ∈ 𝒜^*L*^ (for *L* ≥ *k*) as the average embedding of its constituent *k*-mers, i.e.:

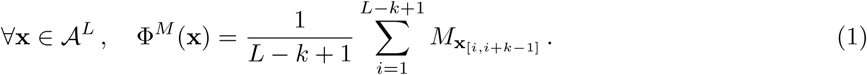

Given the embedding Φ^*M*^ (corresponding to the first layer of the model), we consider a linear model (second layer) of the form:

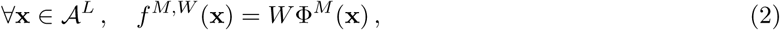

where *W* ∈ ℝ^*T*×*d*^ is a matrix of weights and *T* is the number of taxonomic clades, and the prediction rule:

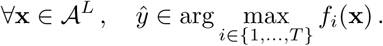

The two-layer classifier is thus parameterized by the two matrices *M* and *W*. Given a training set of reads with known taxonomic clades, which are typically computationally generated by randomly sampling DNA fragments from known organisms and, potentially, adding noise to the fragments, the two matrices are optimized by minimizing an empirical error such as the mean cross-entropy loss over the training set using a standard first-order optimization algorithm such as stochastic gradient descent.

### 2.2 Exploiting *k*-mer symmetry

A limitation of the two-layer classifier is that the size in memory of the representation matrix *M* (with 4^*k*^ ×*d* entries) can quickly become prohibitive as *k*, and the number of *k*-mers, grows. Current implementations of `fastDNA` and similar methods such as DeepMicrobes use for example *k* = 13 and *k* = 12 respectively. On the other hand, it was reported in Vervier *et al.* (2016); Menegaux and Vert (2019) that increasing *k* generally improves the accuracy of the models. We therefore look for strategies to reduce the memory footprint of *M*, in order to explore models with larger *k*’s.

The idea we pursue in this paper is to bin *k*-mers into *N* < 4^*k*^ groups, and to enforce all *k*-mers in any given group to have the same vector representation. Computationally, this allows to reduce the memory footprint of the model by storing only a *N* × *d* matrix of vector representations for the groups, in addition to a lookup table or function to quickly map each *k*-mer to its group.

A first idea to roughly halve the memory footprint of the embedding matrix is to exploit the natural symmetry of DNA, and impose that a read and its reverse complement are indistinguishable in the representation space. The same then goes for *k*-mers: a *k*-mer and its reverse complement should have the same representation. Hence a first step is to learn an embedding per canonical *k*-mer, which roughly halves the number of embeddings. Formally, if for a *k*-mer *x* we denote its reverse complement by 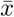, then the canonical *k*-mer 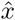 is the smallest of *x* and 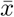 in the lexicographic order. We detail in the Appendix how the lookup mapping from a *k*-mer to its canonical form is performed, and implemented this modification in the vanilla `fastDNA` software.

### 2.3 Exploiting *k*-mers binning with the de Bruijn graph

Exploiting *k*-mer symmetries can roughly halve the memory footprint of the representation matrix *M*, which is not enough to really increase *k* since each increase of *k* by 1 multiplies the number of *k*-mers by 4. Here we propose another, more drastic grouping of *k*-mers by exploiting the idea that *k*-mers that always appear together in the same reads contain the same information for classification purposes, hence should have the same embedding. Indeed, due to the form of embedding for a read (1), one can see that if several *k*-mers are always present in or absent together from a set of reads, then their contribution to the embedding of any read in the set will always be either zero if they are absent from the read, or a constant equal to the sum of their individual embeddings if they are present in the read. Applying this observation to the reads used to train the two-layer classifier, we conclude that the only information that can be used to optimize the *k*-mers representation is at the level of *k*-mers groups. In other words, instead of coding an individual embedding for each *k*-mer, one can get the same modelling capacity by just coding an embedding for that set of *k*-mers, that would correspond to their constant contribution to a read embedding when they are present in the read.

In order to turn this idea into practice, we need a way to bin *k*-mers into groups such that *k*-mers in a group are always present or absent together in the reads used to train the model. For that purpose we exploit the notion of *de Bruijn graph* (dBG), which we now recall. The dBG of a set of sequences *S* is a directed graph (*V, E*) where the vertices V are the *k*-mers appearing in *S*. There is an edge *e* = (*k*_1_, *k*_2_) between two vertices if and only if their corresponding *k*-mers are adjacent in one of the sequences. A maximal non-branching path in a dBG is called a *contig*, or unitig. The compacted dBG (cdBG) merges *k*-mer nodes into contig nodes. Figure 1 illustrates the concepts of dBG and cdBG for a set of two sequences.

**Figure 1:**
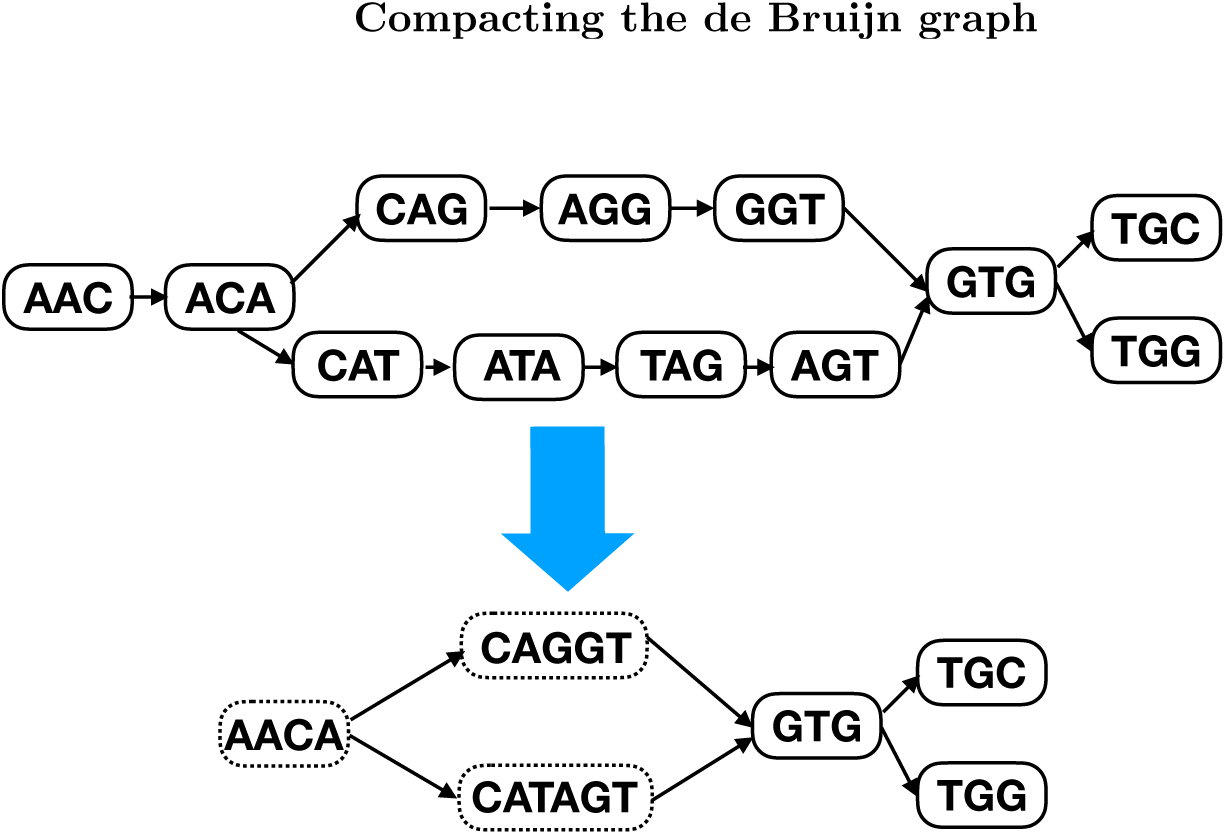
Above: de Bruijn graph for the two sequences AACAGGTGC and AACATAGTGG. Below: corresponding compacted de Bruijn graph. Each node in this graph is a *contig*.

All of the *k*-mers appearing in the reference genomes belong to one and only one contig. As a consequence the number of contigs *N*_*C*_ is bounded by the number of *k*-mers *N*_*k*_.

We now propose to bin *k*-mers by the unique contig they belong to in the cdBG of the training set of reads, and only learn a vector representation for each contig. The embedding of a *k*-mer is then simply the embedding of the contig it belongs to. This reduces the memory footprint of the embedding matrix from 4^*k*^ × *d* to *N*_*C*_ × *d*, which can be a strong reduction when there are many less contigs than possible *k*-mers. For example, in Figure 2 we see that as soon as *k* is larger than 15 then the number of contigs flattens and becomes orders of magnitude smaller than the number of possible *k*-mers, illustrating the potential benefits of the approach.

**Figure 2:**
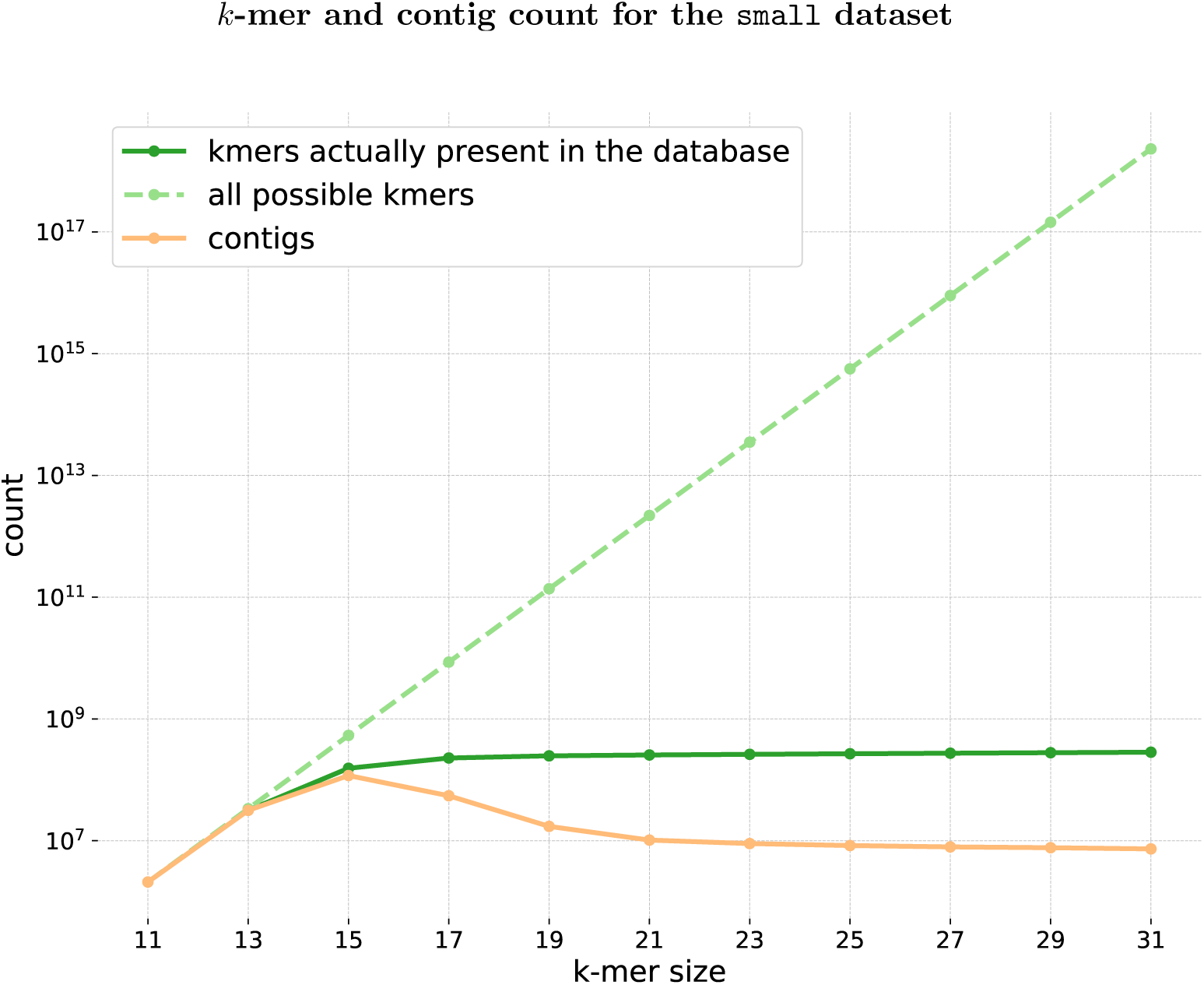
Comparison of the number of *k*-mers and number of contigs present in the reference genomes of the `small` dataset. The dotted line is the theoretical number of possible canonical *k*-mers (= 4^*k*^*/*2)

A caveat with this approach is that *k*-mers that do not appear in the training set of reads have no associated contig. Such *k*-mers occur in a read for which we want to make a prediction, and is even likely to occur more frequently when *k* is large. For example, in Figure 2 we see that for *k* ≥ 15 there is a gap between *k*-mers present in the training set and possible *k*-mers. By default, we ignore *k*-mers absent from the training set by setting their embedding to zero. This also suggests that combining large and small *k*-mers in a single model may be beneficial in some cases, a direction we explore empirically below.

#### Algorithm 1 Brume embedding

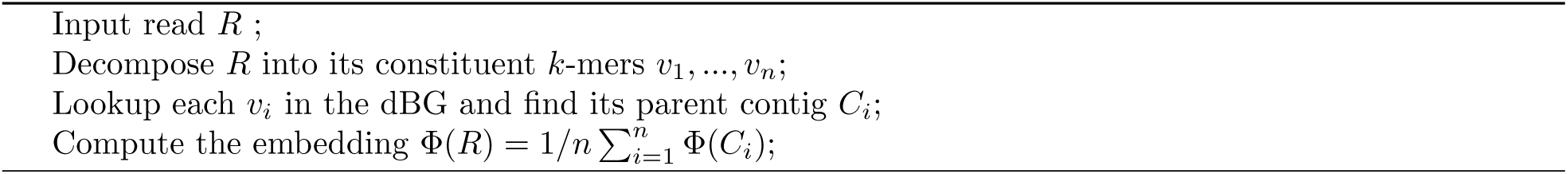

### 2.4 Implementation

We implemented the model as an extension of `fastDNA`. To build the de Bruijn graph, we reused the implementation of `kallisto` (Bray *et al.*, 2016). The source code is freely available and published on github https://github.com/rmenegaux/fastDNA, on the `kallisto` branch, along with scripts to reproduce the presented results.

## 3 Methods

### 3.1 Data

We test our method on the two datasets proposed in Vervier *et al.* (2016) and Menegaux and Vert (2019), called `small` and `large`. Both were extracted from the NCBI bacterial database (reference), and both have a training set and a validation set. In `small` (resp. `large`) the training database is composed of 356 (resp. 2961) bacterial genomes, coming from 51 (resp. 774) species. The validation database is 52 (resp. 193) different genomes, coming from the same 51 (resp. 774) species.

For both training and validation, reads of length *L* = 200 are randomly extracted from the reference genomes. We call a training *epoch* a set of reads such that each base-pair appears once on average (coverage of 1). Epochs are generated on the fly directly during training. We perform data augmentation by adding the possibility to inject random mutations to training reads at a fixed rate *r*, as this was shown to improve performance by Menegaux and Vert (2019)

For validation, we create 4 separate sets from the validation genomes. Each contains about 3.5M (134K for `small`) reads of length *L* = 200, corresponding to a coverage of 1. The first set, called *no-noise*, is extracted as is, with no sequencing noise. The 3 others are extracted with `grinder` Angly *et al.* (2012), a software specifically designed to mimic sequencing noise from real machines. *Balzer* emulates the Roche 454 technology, and is simulated with a homopolymeric error model (Balzer *et al.*, 2010). *Mutation-2* and *mutation-5* are simulated with the 4th degree polynomial proposed by Korbel *et al.* (2009) to study the effect of substitutions, insertions and deletions. The median level of mutations is 2% and 5%, respectively. *Balzer* and *mutation-2* contain error-levels expected from current technology, whereas *mutation-5* is intended to be an extreme case.

We report as metrics the average species-level recall and accuracy, as well as the F1-score to balance between the two.

### 3.2 Reference methods

We compare our results to a standard aligner, BWA-MEM (Li (2013)), that is commonly used in production for metagenomics analysis. We also compare to the standard `fastDNA` method with best parameters from Menegaux and Vert (2019) (*k* = 13, training with mutation noise *r* = 4%), in order to assess the benefits of increasing *k* beyond *k* = 13.

## 4 Results

### 4.1 Embedding canonical *k*-mers

Learning an embedding for each canonical *k*-mer, rather than for all of them, cuts memory usage by roughly a factor of 2. This enables training a `fastDNA` model with *k* = 14 and *d* = 50, rather than the state of the art *k* = 13, *d* = 100 presented in Menegaux and Vert (2019). As shown in Figure 3, this new model is better in both sensitivity and specificity than the previous best, for all levels of sequencing noise, on the `large` dataset. For reads with little noise, it is also better than the alignment method BWA, the latter’s performance being however more robust to sequencing noise. This confirms the findings of Vervier *et al.* (2016); Menegaux and Vert (2019) that increasing *k* in machine learning-based compositional models can be beneficial, and lead to model competitive with alignment-based approaches.

**Figure 3:**
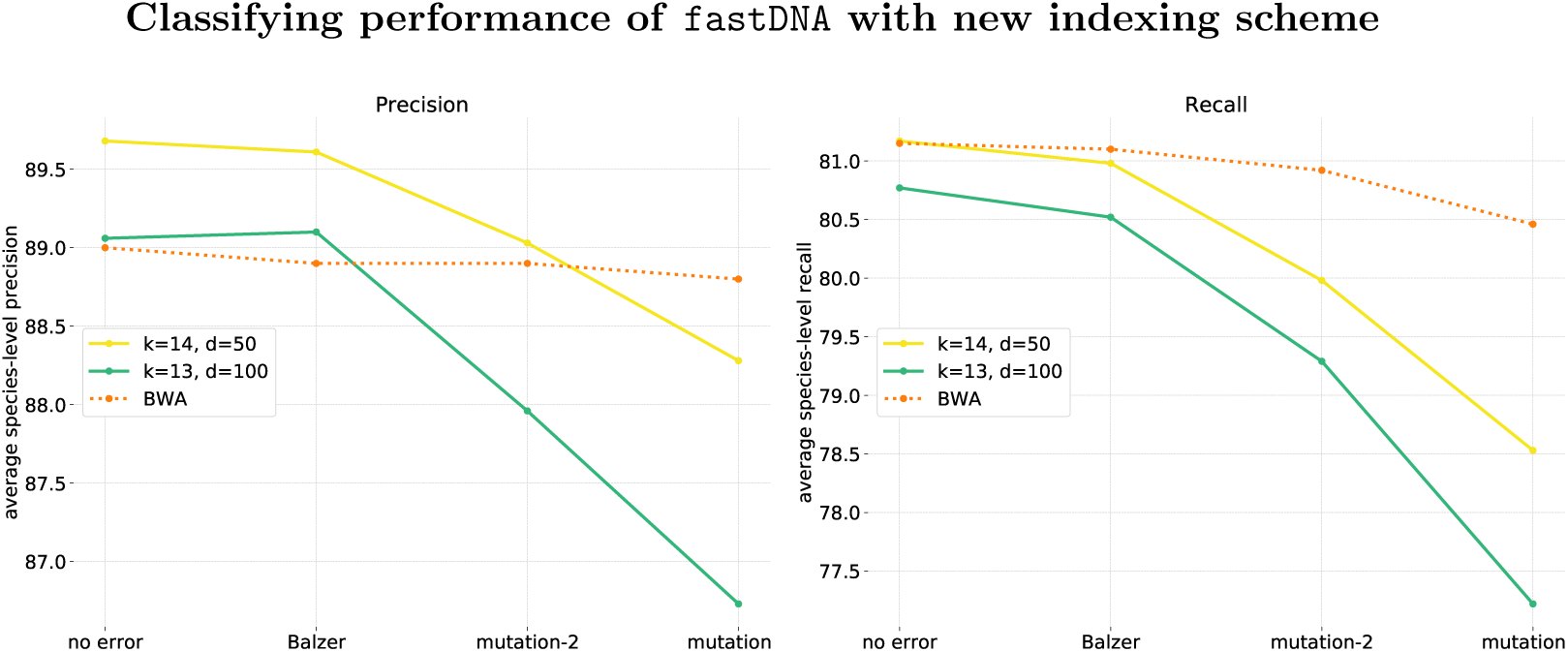
Performance of `fastDNA` with our new indexing (yellow) on the `large` dataset. We compare to the results of the BWA aligner (orange dotted line), and to the state of the art `fastDNA` method, with parameters *k* = 13, dimension *d* = 100 (green). Training mutation rate for both models was *r* = 4%

### 4.2 Embedding contigs

For the rest of this section, we focus on the proposed algorithm `Brume`, which learns an embedding per contig of the cdBG of the training set in order to reduce the memory footprint of the embedding layer. This allows us to explore machine learning-based compositional models for larger values of *k*.

### 4.2.1 Performance on the small dataset

Figure 4 shows the performance of `Brume` on the `small` dataset as a function of *k*, with *k* ranging from 11 to 31. Notice that due to memory constraints, previous work has only investigated values of *k* up to 12 or 13. Notice also that we stopped at *k* = 31 not for memory reasons, but because the performance of the models do not seem to improve by further increasing *k*. Finally, note that for *k* less than 13, there is almost a one to one mapping between *k*-mers and contigs, so results between `fastDNA` and `Brume` are expected to be identical.

**Figure 4:**
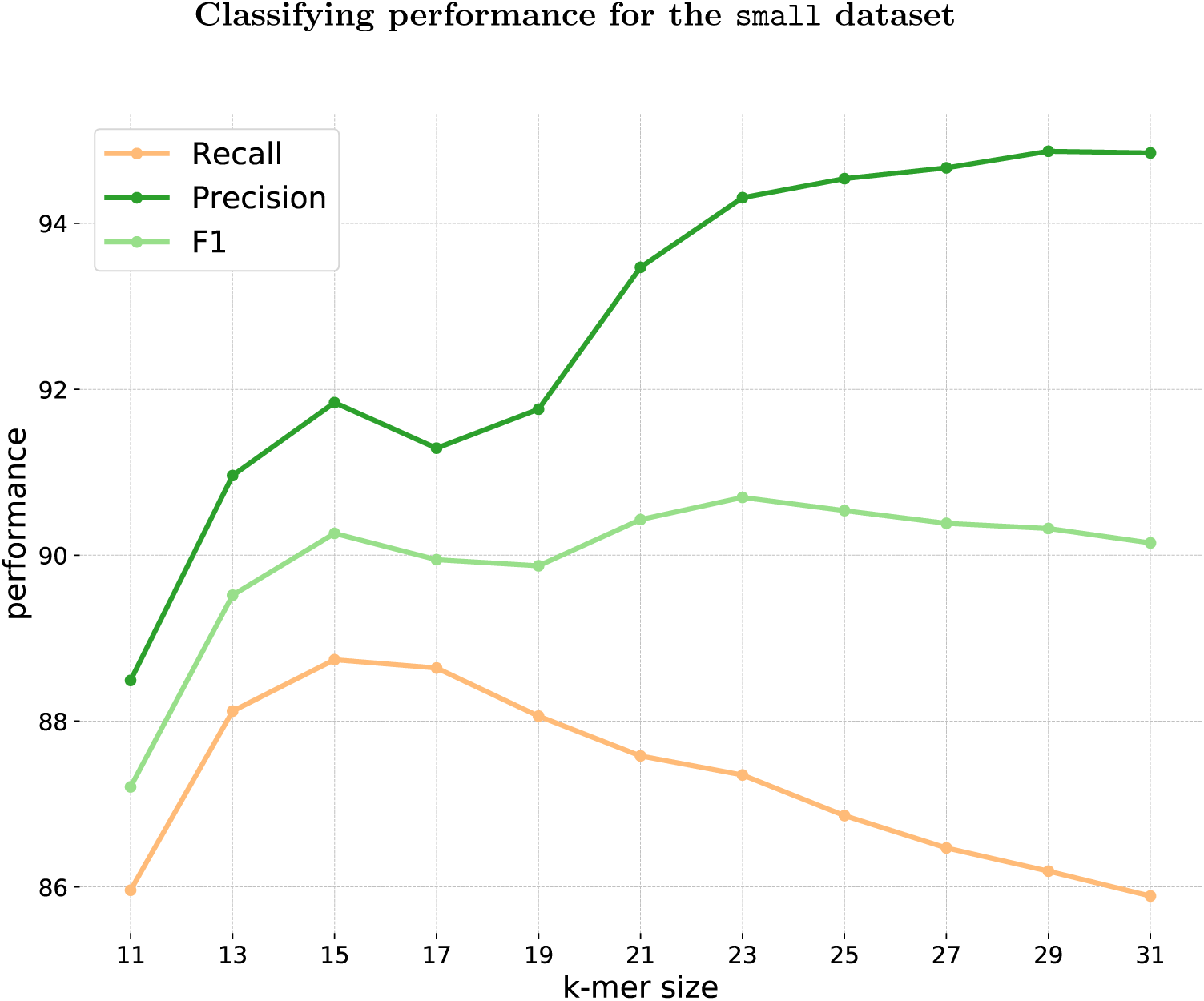
Average species-level precision, recall and F1-score for the `small` dataset, as a function of *k*-mer length. The model were trained for 50 epochs with an embedding dimension of *d* = 10

We see in Figure 4 that recall is maximum at *k* = 15, which coincides with the maximum number of contigs suggesting that perhaps the shattering dimension is what counts ultimately. Larger *k* lead systematically to a decrease in recall, and to an increase in precision from *k* = 17 upwards. The F1-score is optimum for *k* = 23, which shows that our approach is promising.

#### 4.2.2 Performance on the large dataset

We now turn to the more challenging and realistic `large` datasets. Figure 5 shows the performance on the `large` dataset as a function of *k*. As reference we show the performance of BWA, and the best performance of `fastDNA` achieved by embedding canonical *k*-mers (achieved for *k* = 14, *d* = 50, *r* = 4%). We do not have values for *k* = 17 and *k* = 19 because the dBGs were too large to be built or to fit in memory.

**Figure 5:**
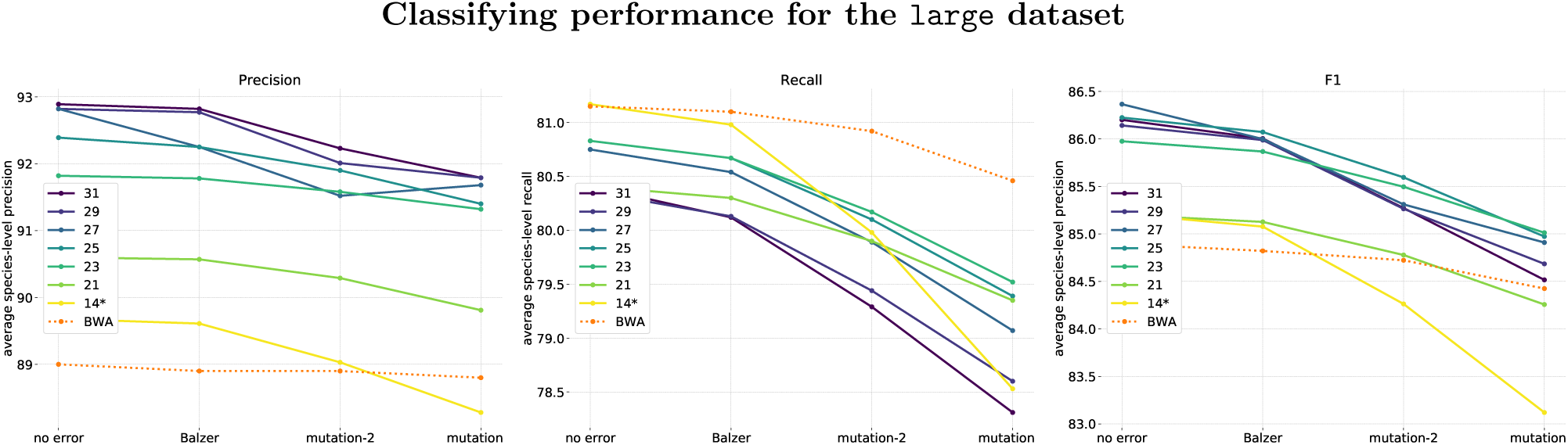
Performance of our method on the `large` dataset, as a function of *k*-mer length *k* (ranging from 21 to 31). The embedding dimension is *d* = 50, and the models were trained for 50 epochs. We compare to the results of the BWA aligner (orange dotted line), and to the `fastDNA` method (yellow), with best parameters *k* = 14 and training mutation rate *r* = 4%)

Overall, we see that, like on the `small` dataset, there is an important benefit in F1 score when *k* increases. In particular, for *k* between 23 and 31, the F1 score of `Brume` is above both `fastDNA` and BWA on all four datasets.

An interesting observation is that models with larger *k*’s look generally more robust to sequencing errors. One can see in Figure 5 that their performance, in terms of F1, recall and precision, decreases more slowly than that of `fastDNA` with higher levels of noise. A possible explanation is that one mutation in a read leads all *k* neighboring *k*-mers to be corrupted. The larger *k* is, the more specific they are and most of them will not find a match in the cdBG, and therefore will not impact the embedding or the classification. On the contrary, `fastDNA` treats these erroneous *k*-mers as any other.

Similarly to the *small* dataset, the increase in F1 score for larger *k*’s hides the fact that recall (resp. precision) consistently decreases (resp. increases) with larger *k*’s. The increase in precision for larger *k* can be at least partly attributed to an increased proportion of reads that are not classified. As seen in figure 6, this proportion can go over 10% for the `large` dataset, and increases with both *k* and the level of noise. Unclassified reads are reads with no matching *k*-mers found in the dBG of the training set, and the predictions are left blank for them: they count as errors for recall but do not count for precision. The fact that precision goes up as recall goes down suggests that those missing reads are precisely those which are difficult to classify.

**Figure 6:**
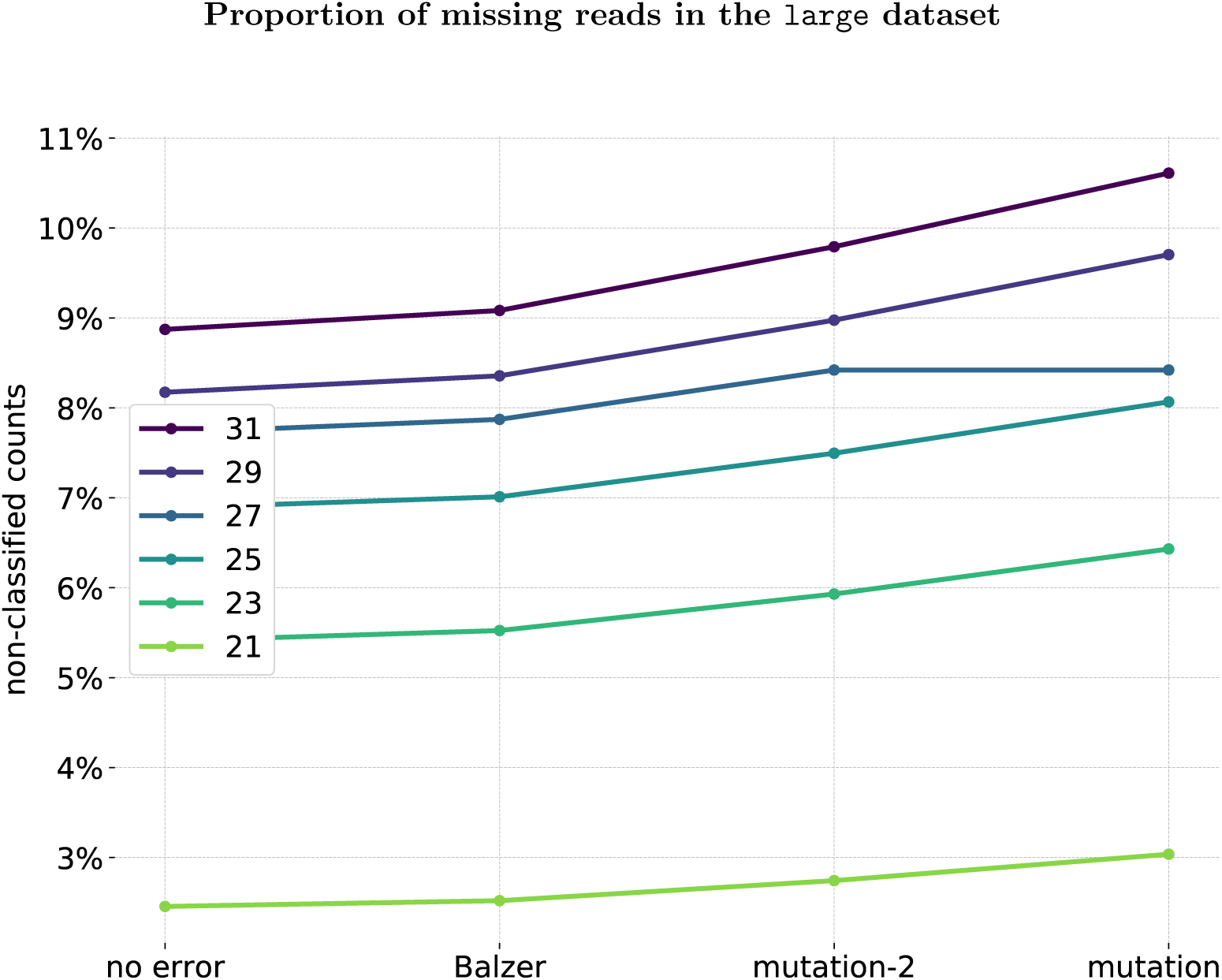
Proportion of validation reads in `large` with no matches in the reference genomes as a function of simulated sequencing noise. Each line is a fixed value of *k*-mer length *k*.

## 5 Discussion

We demonstrated that *k*-mer-based machine learning methods for read classification can be extended to large *k*, by quantizing the *k*-mer space. We did so by using the de Bruijn graph and giving the same representation to *k*-mers that were in the same contig. A supervised classifier, similar in architecture to `fastDNA`, outperforms state of the art in terms of precision and F1-score.

When *k* gets large, *k*-mers become increasingly specific to each read and we are increasingly confronted to the situation where a read to be classified has no *k*-mer in common with the training set used to train the model. In that situation, we do not classify the read, which decreases the recall of our model, but as we observed also increases the precision since those reads tend to be “hard” to classify. In order to try to classify reads that have been left out, we could also make use of smaller, dense *k*-mers to give them a representation. This naturally leads to the idea of mixing representations with large and small *k*. We have tried several ways to do so, in particular:

1. Training a separate `fastDNA` model with short *k* that we use only in case `Brume` cannot match the read
2. Training separate embeddings for short *k* and long *K*, then averaging or concatenating them to yield a unified embedding.

We implemented and tested these methods, but overall the classification performance on the homeless reads is not good enough to justify the extra workload. For example, in Figure 7, we show the results for a hybrid model *k* = 13, *K* = 31, which gives a *d*-dimensional embedding to both short *k*-mers and contigs, then averages the two to produce the final representation of a read. Recall is indeed boosted but at a high cost in precision.

**Figure 7:**
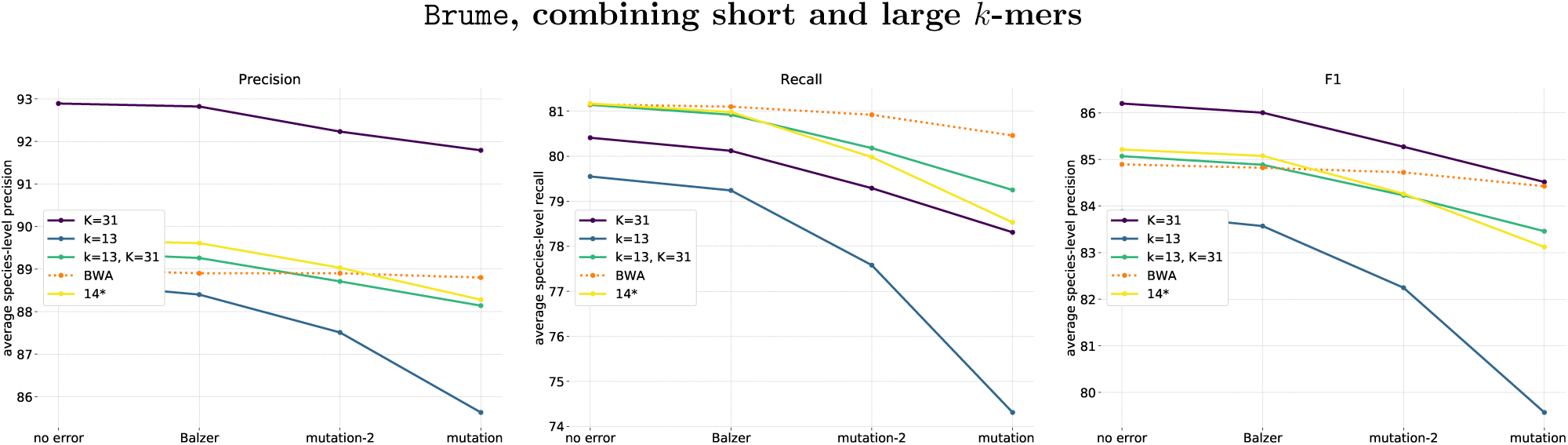
Performance on the `large` dataset of a hybrid model. The embedding of a read is the average of the embeddings of its 13-mers and its 31-mers. Dimension is *d* = 50. Shown as comparison are `Brume` with *k* = 31 and `fastDNA` with *k* = 13.

This shows that combining short and long *k*-mers as we did is not enough to improve both precision and recall, and suggests that increasing *k* allows to navigate a trade-off between precision and recall, where models with larger *k*’s simply do not classify “hard” reads and are more accurate on “easy” reads.

Regarding speed, the *k*-mer lookup in the dBG comes with a significant impact on classification speed. `kallisto` have a heuristic based on the dBG to avoid looking up every *k*-mer, which we have not reimplemented as of now. We have also experimented with another library to build the dBG called `bifrost` (Holley, 2019), which is slightly faster, presumably because of the rolling hash they use to lookup *k*-mers. The built dBGs were essentially the same and therefore the classifying performance of `Brume` was unchanged. Results are not shown here.

## 6 Conclusion

We have proposed a new way to bin *k*-mer representation for machine learning-based models, using a binning based on contigs in a cdBG, and illustrated the benefits of using larger *k* values than currently available on a metagenomics application. We believe these embeddings can be used for other more sophisticated models, such as deep learning-based models, and in other fields than metagenomics, such as RNA-seq.

## Appendix Indexing scheme

Because of the natural symmetry of DNA, a read and its reverse complement should be indistinguishable in our model. The same goes for *k*-mers: a *k*-mer and its reverse complement should have the same representation.

Let *M*^*k*^ = {(*u, ū*), *u* ∈ 𝒜^*k*^, *u* ≤ *ū*} the set of alphabetically ordered pairs of *k*-mers and their reverse complements. Let *m*_*k*_ = ‖*M*^*k*^‖. A palindrome is a *k*-mer *u* such that *u* = *ū* The number of elements in *M*^*k*^ is half the number of elements in 𝒜^*k*^ plus half the number of palindromes, so *m*_*k*_ = 4^*k*^*/*2 if *k* is odd (no palindromes) and *m*_*k*_ = 4^*k*^*/*2 + 2^*k*^ if *k* is even.

An indexing scheme is a bijection *ϕ* from *M*^*k*^ to [0, *m*_*k*_ − 1].

We define *ϕ* recursively on *k* as follows:

1. *ϕ* of the empty string is 0: *ϕ*(″) = 0
2. *ϕ* of a single character *a* ∈ 𝒜 is *ϕ*(*a*) = {`A`: 0, `C`: 1, `G`: 2, `T`: 3}[*a*]:
3. For *u* ∈ 𝒜^*k*^, we write *u* = *u*_1_*u*′*u*_*k*_, where *u*_1_, *u*_*k*_ ∈ 𝒜 are the first and last characters of *u*, then use the rule described in Algorithm 2

### Algorithm 2 fastDNA symmetric and bijective indexing scheme

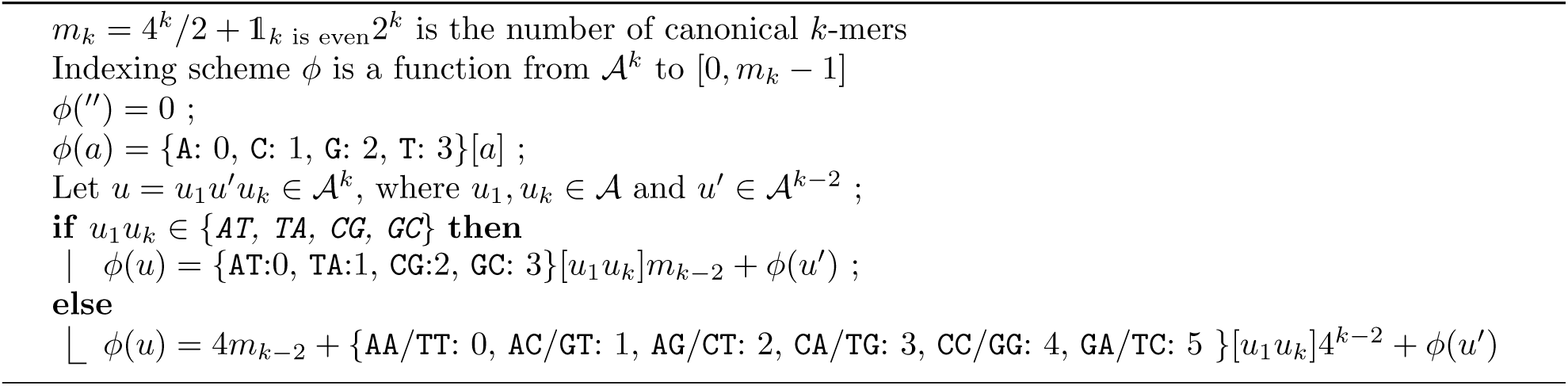

